# Functional and long-lived melanocytes from human pluripotent stem cells with transient ectopic expression of JMJD3

**DOI:** 10.1101/2023.03.03.530736

**Authors:** Chie Kobori, Ryo Takagi, Ryo Yokomizo, Sakie Yoshihara, Mai Mori, Hiroto Takahashi, Palaksha Kanive Javaregowda, Tomohiko Akiyama, Minoru S.H. Ko, Kazuo Kishi, Akihiro Umezawa

## Abstract

**Background:** Melanocytes are an essential part of the epidermis, and their regeneration has received much attention because propagation of human adult melanocytes *in vitro* is too slow for clinical use. Differentiation from human pluripotent stem cells to melanocytes has been reported, but the protocols to produce them require multiple and complex differentiation steps.

**Method:** We differentiated human embryonic stem cells (hESCs) that transiently express JMJD3 to pigmented cells. We investigated whether the pigmented cells have melanocytic characteristics and functions by qRT-PCR, immunocytochemical analysis and flow cytometry. We also investigated their biocompatibility by injecting the cells into immunodeficient mice for clinical use.

**Result:** We successfully differentiated and established a pure culture of melanocytes. The melanocytes maintained their growth rate for a long time, approximately 200 days, and were functional. They exhibited melanogenesis and transfer of melanin to peripheral keratinocytes. Moreover, melanocytes simulated the developmental processes from melanoblasts to melanocytes. The melanocytes had high engraftability and biocompatibility in the immunodeficient mice.

**Conclusion:** The robust generation of functional and long-lived melanocytes are key to developing clinical applications for the treatment of pigmentary skin disorders.

## Background

Skin is a vital organ that maintains body fluid and temperature balance against external environments. Although melanocytes are small population cells in the epidermis, they play an important role in protecting skin from harmful ultraviolet rays. Vitiligo, the most common pigmentary disorder, affects approximately 0.1 to 2.0% of the world’s population (1), and causes severe psychological distress, especially when it involves the exposed skin, such as on the face or hands, especially when patients have dark skin (1-3). The first choice is medical therapies such as psoralen, ultra-violet A, and immunomodulating therapy. Treatment is less successful if vitiliginous lesions are located on the back of the hands, eyelids, or around the mouth, which affects strikingly their appearance and mainly requires treatment. In these cases, surgical therapy is chosen: Transplantation of autologous melanocytes has been investigated for efficacy (4). However, the proliferation rate of melanocytes *in vitro* depends on the donor site and age. Melanocytes from the newborn foreskin grow the fastest, while those from the arm of adults are the slowest (5). Obtaining the necessary quantity of melanocytes for transplantation is difficult because the therapies are often performed in adults.

To solve this problem, the generation of human melanocytes from pluripotent stem cells was introduced first in 2006 (6), and several strategies have been published thereafter (7-9). The differentiation protocol should be simple so that the manufacturing process will be more robust, validated and reliable. The aim is to make this technology practical for future clinical applications because the demand for vitiligo treatment is high.

We have prepared bio-artificial organs with functional cells in humans (10, 11). Pigmented cells appeared with a certain probability during hepatic differentiation of human pluripotent stem cells. In this study, we reproducibly generated pigmented cells with the same life span, properties, and functions as human tissue-derived melanocytes. The simple protocol described in this study will realize future clinical applications since it does not require complex tuning of media or reagents.

## Methods

### Melanocytic differentiation

We used an hESC line that transiently expresses the C-terminal region (catalytic domain) of histone demethylase JMJD3 in a doxycycline (DOX, Takara Bio Inc, Japan) inducible manner (12, 13). The hESC line was cultured on a feeder layer of freshly plated gamma-irradiated mouse embryonic fibroblasts (MEFs) isolated from ICR embryos at 12.5 d gestation and passaged two times before irradiation (30 Gy), in the ESC culture media [KNOCKOUT-Dulbecco’s modified Eagle’s medium (KO-DMEM) supplemented with 20% KNOCK-OUT-Serum Replacement (KO-SR), 2 mM Glutamax-I, 0.1 mM non-essential amino acids (NEAA), 50 U/ml penicillin-50 μg/ml streptomycin (Pen-Strep), 0.055 mM β-mercaptoethanol and 10 ng/ml recombinant human full-length bFGF (All reagents from Gibco, USA)].

Before embryoid body generation, 2 μg/ml of DOX was added to the hESC culture media for two days to overexpress JMJD3. One day apart, hESCs were dissociated into single cells and cultivated in the 96-well plates in the XF32 medium [85% KNOCKOUT DMEM, 15% KNOCKOUT Serum Replacement XF CTS (Life Technologies), 2 mM GlutaMAX-I, 0.1 mM NEAA, Pen-Strep, 50 μg/mL l-ascorbic acid 2-phosphate (Sigma-Aldrich, USA), 10 ng/mL heregulin-1β (recombinant human NRG-β 1/HRG-β1 EGF domain; R&D Systems, USA), 200 ng/mL recombinant human IGF-1 (LONG R3-IGF-1; Sigma-Aldrich, USA), and 20 ng/ mL human bFGF (Life Technologies)] with 10 μM Rho kinase inhibitor Y-27632 (Wako, Japan) for 6 days. The EBs were transferred to the 24-well plates coated with 30 μg/cm^2^ collagen type I and cultivated in the XF32 medium for five days. The cells were further cultivated and propagated in XF32 medium without bFGF for ten days and then in ESTEM-HE medium (Emukk co., Japan) for 15 days at 37°C in a humidified atmosphere containing 95% air and 5% CO2. In the middle of that time period, as antibiotic selection, 1 μg/ml of puromycin (Takara) was added to the medium only for two days.

### Purification and proliferation of melanocytes

When the cultures reached subconfluence, the cells were dissociated with Trypsin–EDTA solution (IBL CO., Ltd, Japan), and seeded at a density of appropriately 5 × 10^5^ cells in a 100-mm dish. Medium changes were carried out three times a week after that. The cells were co-cultured with MEFs at the first passage, and then on dishes coated with 0.25 μg/cm^2^ LN-511-E8 (Nippi, Japan) alone after the second passage. The medium was ESTEM-HE medium during purification and changed to Human Melanocyte Basal Medium Kit (Gibco) or Melanocyte Growth Basal Medium-4 (Lonza, Switzerland) after pigmented cells were dominant.

### Co-culture of Melanocytes and Keratinocytes

Co-culture of melanocytes and keratinocytes was performed to investigate melanocyte function, i.e. transportation of melanin to keratinocytes. Human postnatal epidermal melanocytes were seeded onto culture dishes coated with 0.25 μg/cm^2^ LN-511-E8 at a density of 2.0 × 10^4^ cells/cm^2^. Two days later, mouse feeder cells were seeded at a density of 2.0 × 10^4^ cells/cm^2^ (8). Briefly, the feeder cells treated with 10 μg/ml mitomycin C (Nacalai tesque, Japan) for 2 hours were seeded at 2.0 × 10^4^ cells/cm^2^ in α-MEM (Thermo Fisher Scientific) containing 10% FBS (Thermo Fisher Scientific), 100 unit/mL Pen-Strep and 0.25 μg/mL Fangizon (Bristol Myers Squibb, Japan). The next day, human postnatal epidermal keratinocytes were seeded at a density of 1.0 × 10^4^ cells/cm^2^ in the keratinocyte culture medium (8).

### Quantitative reverse transcriptase-PCR (qRT-PCR)

Total RNA was prepared by using ISOGEN (Nippon Gene, Japan) and PCR Inhibitor Removal Kit (Zymo Research, USA). The RNA was reverse transcribed to cDNA by using Superscript III Reverse Transcriptase (Invitrogen, USA) with ProFlex PCR System (Applied Biosystems, USA). qRT-PCR was performed on QuantStudio 12 K Flex (Applied Biosystems, USA) using a Platinum SYBR Green qPCR SuperMix-UDG (Invitrogen, USA). Expression levels were normalized with the reference gene, glyceraldehyde-3-phosphate dehydrogenase (GAPDH). The primer sequences are shown in Table 1.

**Table 1.**
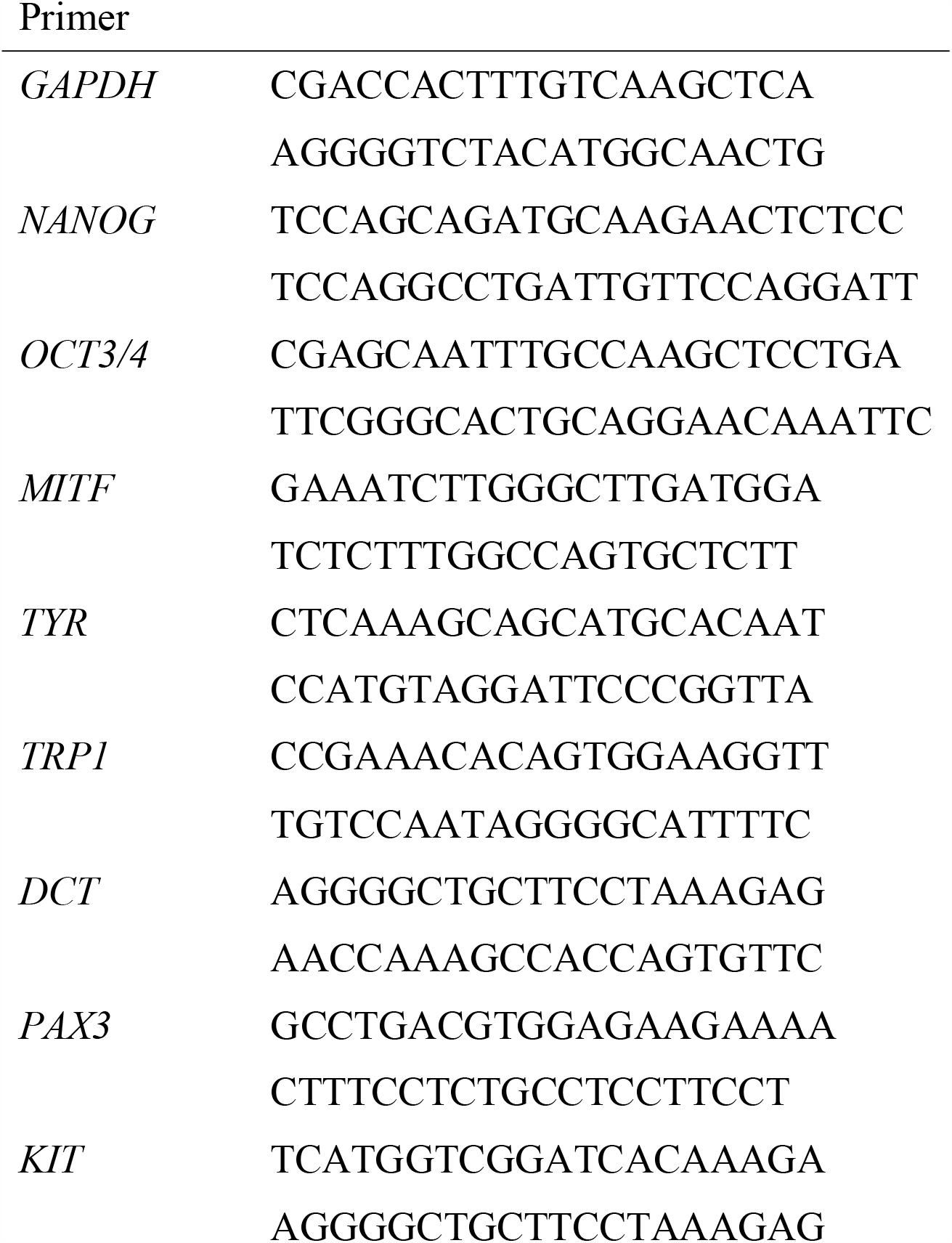
Primer List.

### Immunocytochemical Analysis

Cells were fixed with 4% paraformaldehyde for 10 minutes at room temperature. After being washed with PBS and treated with 0.1% Triton X-100 (Nacalai tesque) for 10 minutes, they were exposed to Protein Block Serum-Free Ready-To-Use (DAKO, Denmark) for 30 minutes at room temperature, and then incubated overnight at 4°C in primary antibodies diluted with 1% BSA. Following washing with PBS, they were incubated for 30 minutes at 4°C in secondary antibodies (1:500 diluted with BSA) and DAPI (1:1000 with BSA). Then, they were mounted with Fluorescence Mounting Medium (DAKO). Antibody information is provided in Table 2.

**Table 2.**
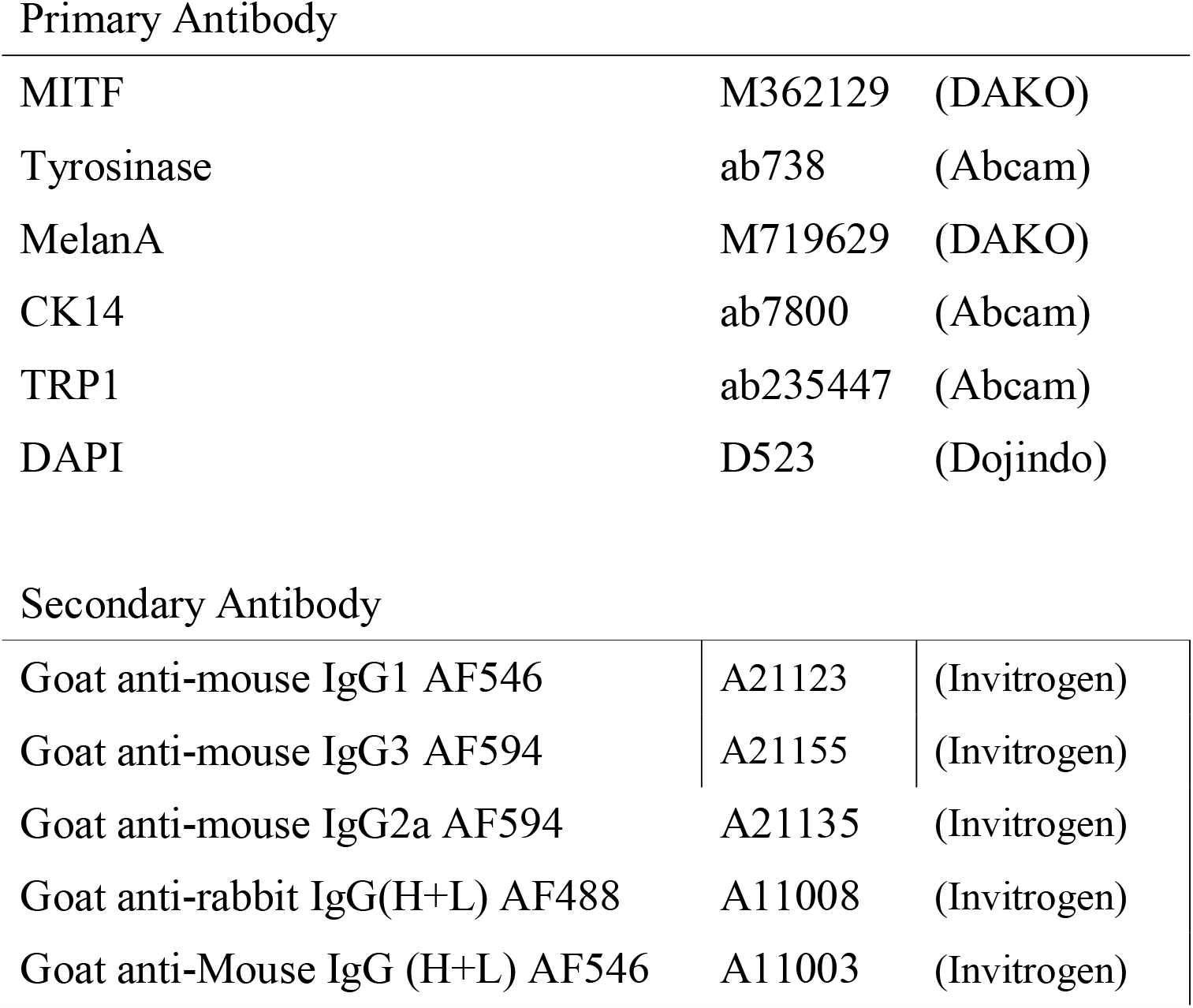
Antibody List.

Primary Antibody
|  |  |  |
| --- | --- | --- |
| MITF | M362129 | (DAKO) |
| Tyrosinase | ab738 | (Abcam) |
| MelanA | M719629 | (DAKO) |
| CK14 | ab7800 | (Abcam) |
| TRP1 | ab235447 | (Abcam) |
| DAPI | D523 | (Dojindo) |
| Secondary Antibody |  |  |
| Goat anti-mouse IgG1 AF546 | A21123 | (Invitrogen) |
| Goat anti-mouse IgG3 AF594 | A21155 | (Invitrogen) |
| Goat anti-mouse IgG2a AF594 | A21135 | (Invitrogen) |
| Goat anti-rabbit IgG(H+L) AF488 | A11008 | (Invitrogen) |
| Goat anti-Mouse IgG (H+L) AF546 | A11003 | (Invitrogen) |

### Flow cytometry

Cells were fixed and permeabilized with Fixation/Permeablization Solution Kit (BD Cytofix/Cytoperm). Briefly, cells were exposed to Fixation/ Permeabilization solution for 20 minutes at 4°C and washed with 1x BD Perm/Wash™ buffer. The cells were then incubated for 30 minutes at room temperature in primary antibodies diluted with 3% FBS/PBS. After washing with FBS/PBS, they were incubated for 30 minutes at room temperature in secondary antibodies (1:200 with FBS/PBS). Negative controls were performed by omitting the primary antibody. Measurement was carried out using Cell Sorter LE-SH800 series (SONY, Tokyo). Antibody information is provided in Table 2.

### Mouse injection

Melanocytes (1 × 10^5^) in 100 μL isotonic sodium chloride solution were subcutaneously injected into two sites of SCID beige mice. The skin was examined at one week and seven weeks after the injection.

## Results

### Pigmented cells were differentiated from human embryonic stem cells

We used hESCs (SEES3) which conditionally overexpressed JMJD3 for melanocytic differentiation. hESCs were induced to express JMJD3 with exposure to DOX and then generated EBs. The EBs were transferred to the 24-well plates coated with collagen type I and cultivated for 15 days. The cells were further cultivated and propagated (Figure 1a). Pigmented cells with dendritic shape appeared 7-10 days after exposure to puromycin. Pigmented cells appeared in all culture plates (Figure 1b). Pigmented cells increased and became dominant in the dish (Figure 1c). Pigmented cells reached nearly 100% at passage 15 (Figure 1d). Pigmented cells proliferated as visible black colonies and the cell pellet was black in color without any stain (Figure 1e).

**Figure 1.**
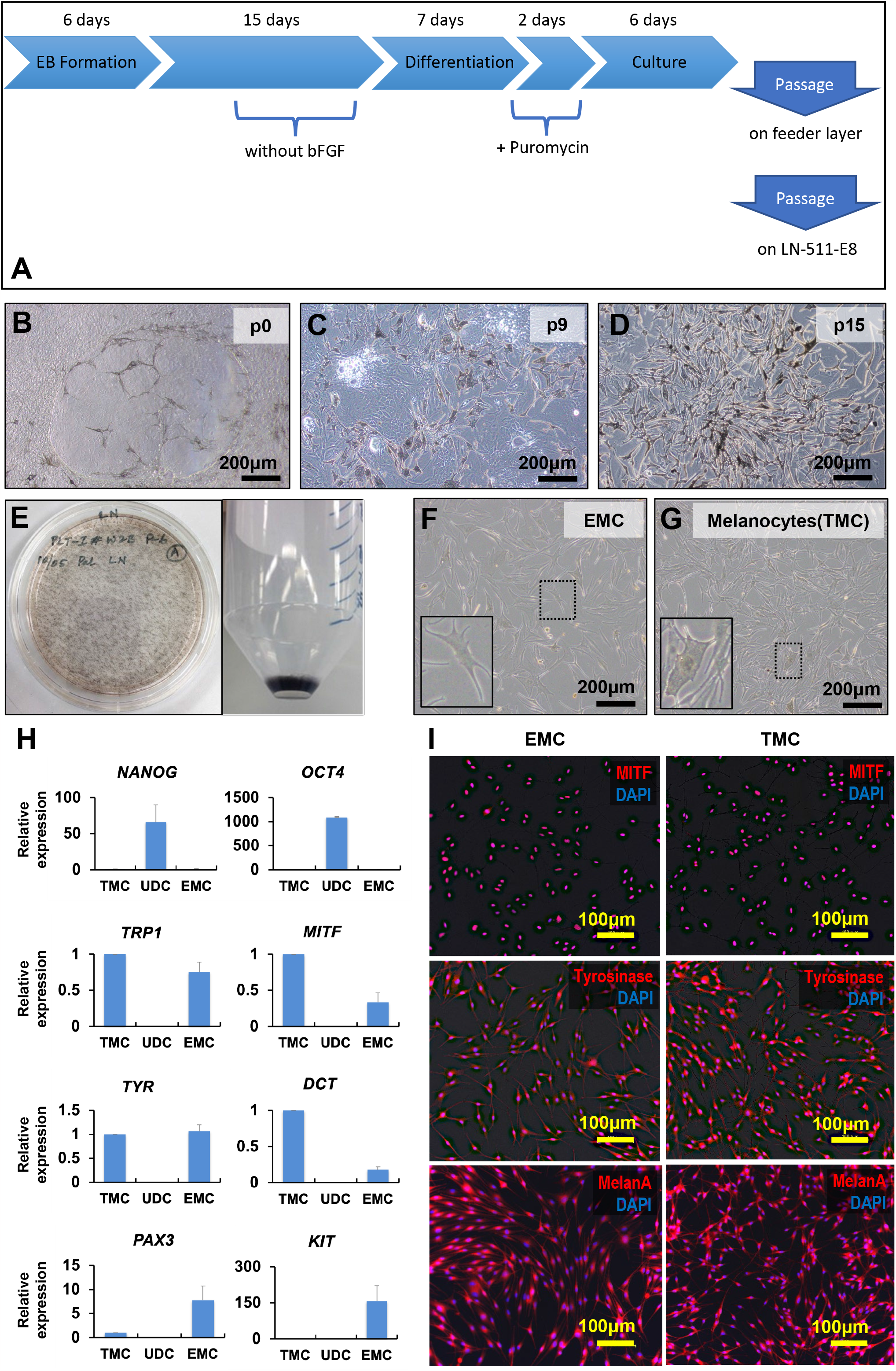
hESC-derived pigmented cells exhibit a melanocytic phenotype. **A**. Graphical illustration of differentiating from embryonic stem cells to melanocytes **B**. Pigmented and dendritic cells appeared 7-10 days after exposure to puromycin at passage 0 (p0). **C**. Pigmented cells increased, while non-pigmented cells decreased around passage 9 (p9). **D**. Pigmented cells were dominant by passage 15. **E**. Gross appearance of pigmented cells. The culture plate was covered with black colonies (left), and the cell pellet was black without any staining (right). **F-G**. Human Embryonic stem cells-derived melanocytes (F: EMCs) at passage 25 resembled tissue-separated melanocytes (G: TMCs) in morphology. **H**. Quantitative RT-PCR analysis of the genes for NANOG, OCT4, TYRP1, MITF, Tyrosinase, PAX3, Kit, and DCT, in undifferentiated cells (UDCs), EMCs, and TMCs. The expression levels were normalized by expressions of glyceraldehyde-3-phosphate dehydrogenase (GAPDH). The expression levels of TMC were set at 1.0. Each expression level was calculated from the results of triplicate technical experiments and the charts are drawn as the average ± standard deviation. **I**. Immunocytochemical analysis of EMCs(left) and TMC (right) with antibodies against MITF (red), Tyrosinase (red), and MelanA (red). Nuclei were counterstained with DAPI (blue).

### Pigmented cells had melanocytic characteristics

We examined the pigmented cells to investigate whether they have melanocytic characteristics. We first compared their morphology with tissue-derived melanocytes (TMCs) (Figure 1f-g). The pigmented cells exhibited multidendritic, star-like morphology with pigmented melanosomes and translucent cytoplasm like TMCs. We then performed qRT-PCR to investigate the gene expression of melanocytic markers in undifferentiated hESCs (UDCs), pigmented cells, and TMCs (Figure 1h). The pigmented cells at passage 15 decreased expression of the genes for *NANOG* and *OCT4*, compared with UDCs, and increased melanocytic markers, i.e. *DCT, TYRP1, MITF*, and *TYR*. The expression levels of melanocytic markers in the pigmented cells were comparable with that in TMCs. It is noteworthy that *PAX3* and *KIT* which are expressed in melanoblasts or premature melanocytes were detected in the pigmented cells but not in TMCs.

Immunocytochemical analysis showed that every pigmented cell had melanocyte-specific proteins, MITF, tyrosinase, and MelanA. Tyrosinase and MelanA were stained in the cytoplasm and MITF was stained in the nucleus (Figure 1i). Taken together, these results indicate that the pigmented cells derived from hESCs were indeed melanocytes. We call the pigmented cells as EMCs (human Embryonic stem cell-derived melanocytes) below.

### EMCs maintained melanocytic characteristics over time

We performed flow cytometric analysis to investigate the proportion of melanocytes in EMCs and TMCs using an antibody against MelanA. Nearly 100% of EMCs at passage 16 were positive for MelanA (Figure 2a). Next, we investigated EMCs’ growth pattern and change in gene expression by qRT-PCR. EMCs and TMCs showed the same growth rate: Doubling time of EMCs and TMCs was almost the same (Figure 2b). EMCs maintained their proliferation rate and differentiated phenotypes by 14 population doublings (passage 33) at 200 days (Figure 2c). The melanoblast markers, *DCT, PAX3*, and *KIT* increased between passage one and passage 21. *MITF* and *TRP1* showed the peak of expression at passage 15 when EMCs were dominant in the culture plate and gradually decreased as the passage proceeded. In contrast, *TYR* increased until passage 32. EMCs seemed to reach replicative senescence at passage 45.

**Figure 2.**
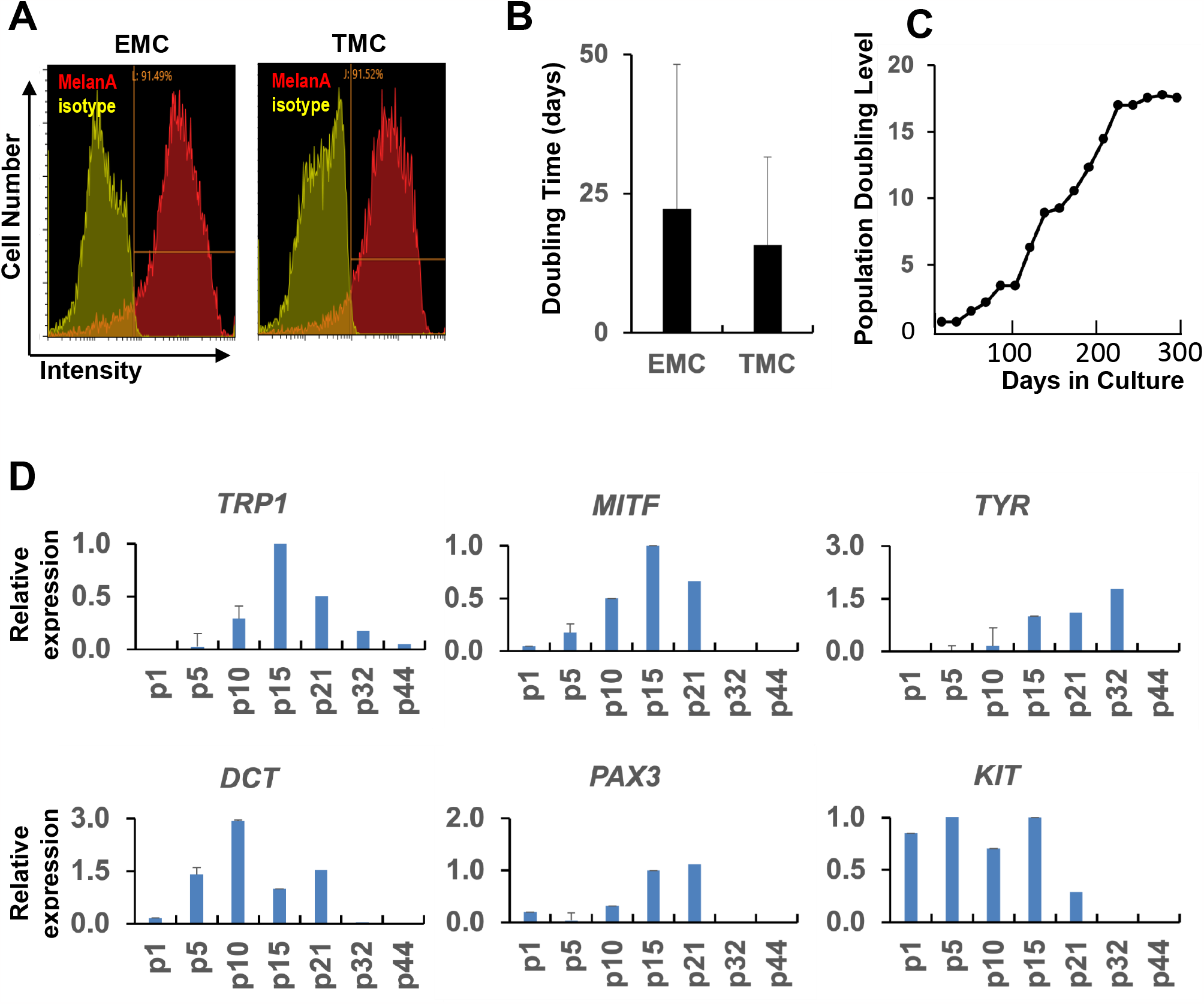
EMCs maintain melanocytic phenotypes at 14 population doublings. **A**. Flowcytometric analysis of EMCs and TMCs with antibodies against MelanA (red). Positive cells of EMCs were 91.49%, while TMCs were 91.52%. **B**. Population doubling level of EMCs from the 20th to 38th passage. “Population doubling” indicates the cumulative number of divisions of the cell population. **C**. Comparison of Doubling Time with EMCs (passage from 20 to 30) and TMC (passage from 2 to 12). The doubling time averages were 22.3 days (±11.45) and 14.2 days (±15.20) in EMCs and TMCs, respectively. **D**. Quantitative RT-PCR analysis of genes for MITF, Tyrosinase, TRP1, DCT, PAX3, and KIT at each passage. EMCs reached 14 population doublings at passage 21 (p21). The expression levels were normalized by expressions of GAPDH. The expression levels of EMC at passage 15 (p15) were regarded as equal to 1.0. Each expression level was calculated from the results of triplicate technical experiments and the charts are drawn as averages ± standard deviation.

### EMCs were capable of transferring melanin to adjacent keratinocytes

We investigated the melanogenesis of EMCs and found that EMCs produced melanin from DOPA as revealed with a DOPA reaction test (Figure 3a). Melanocytes’ function is not only melanin production but also transferring to peripheral keratinocytes. Keratinocytes were co-cultured with EMCs or TMCs for four days and analyzed by immunocytochemical staining. Keratinocytes showed TRP1-positive inclusion in CK14-positive cytoplasm, while keratinocytes alone had no TRP1-positive inclusion (Figure 2b). The transfer of melanin was also analyzed quantitatively by using flow cytometry with antibodies against CK14 and TRP1. CK14+/TRP1+ double-positive keratinocytes were not detected just after both cells were mixed (Figure 3C, 0.27%); However, CK14+/TRP1+ double-positive keratinocytes increased when keratinocytes were co-cultured with EMCs (Figure 3D, left panel, 1.14%). CK14+/TRP1+ keratinocytes with flow cytometry could be keratinocytes with TRP1-positive inclusion with immunocytochemistry (Figure 3B, middle panel). These results suggest that EMCs are capable of not only producing melanin but also transmitting melanin to keratinocytes.

**Figure 3.**
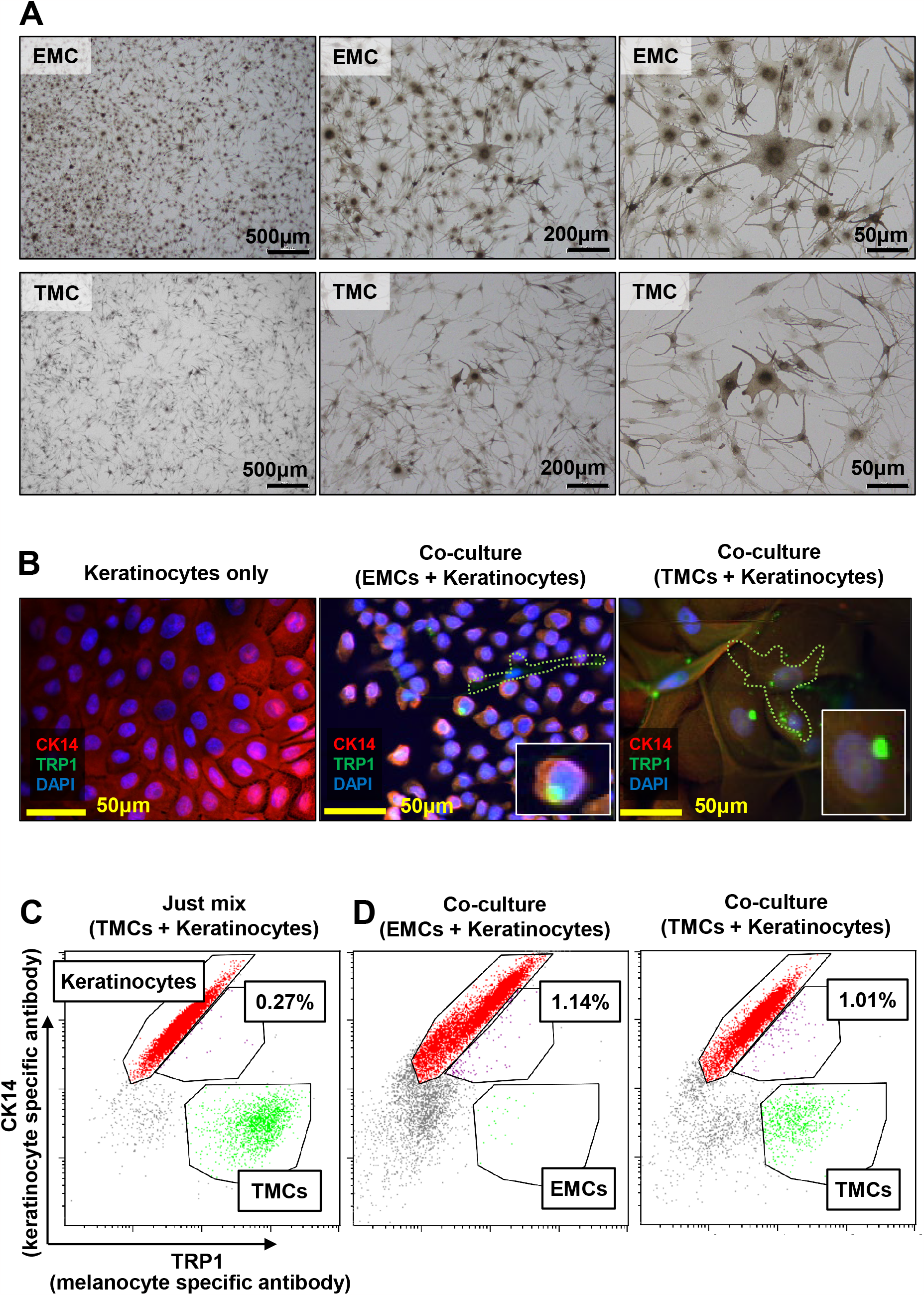
EMCs transfer pigments to keratinocytes in co-culture. **A**. DOPA (3,4-dihydroxyphenylalanine) reaction of EMCs (upper panels) and TMCs (lower panels). **B**. Immunocytochemical analysis of keratinocytes on EMCs or TMCs with antibodies against CK14 (red) and TRP1 (green). Nuclei were counterstained with DAPI (blue). Lower right inlets show magnified keratinocytes with TRP1-positive inclusion. Green dotted lines show the outline of melanocytes. **C-D**. Flow cytometric analysis of EMCs, TMCs, and keratinocytes with antibodies against CK14 and TRP1. Red and green dots indicate CK14+/TRP1-keratinocytes and CK14-/TRP1+ melanocytes, respectively. Purple dots indicate double-positive CK14+/TRP1+ keratinocytes. (C) Keratinocytes and melanocytes were mixed just before the analysis. (D) Keratinocytes were co-cultured with EMCs or TMCs for six days.

### EMCs engraft in vivo over time

We investigated EMC’s engraftability and biocompatibility. EMCs were injected into the back of three mice (Figure 4a). EMCs remained at seven weeks, and no inflammation or tumorigenesis was observed at the injected sites (Figure 4b-c). EMCs did not change in size and position at 1 and 7 weeks and did not modify the macroscopic color of surroundings including skin and hair. (Figure 4d-f). Interestingly, EMCs retained their size and color better than the TMCs that served as a positive control.

**Figure 4.**
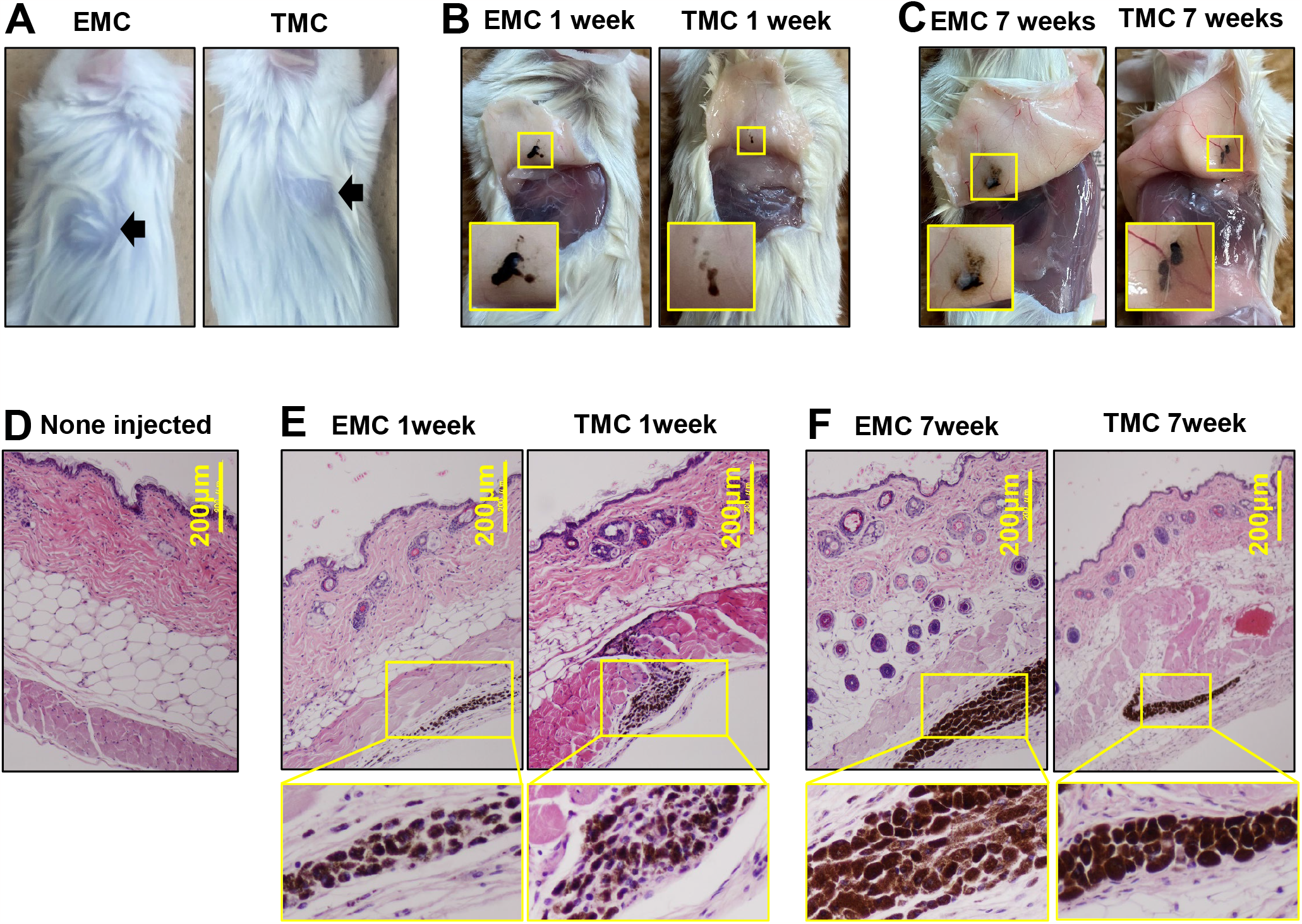
Engraftability and biocompatibility of EMCs. **A**. Gross appearance of the mice just after subcutaneous injection of EMCs and TMCs (arrows). **B-C**. Gross appearance of EMCs and TMCs with pigmentation at one week (B) and seven weeks (C). Lower left inlets show a magnified pigmentation area. **D-F**. Histological analysis of EMC- (E), and TMC- (F) injected skin by H.E. stain. Lower panels show high-power views of EMCs and TMCs.

## Discussion

hESC-derived melanocytes exhibited a high proliferative ability over time with functionality and had the same gene expression pattern during melanocytic development in embryogenesis. Melanocytes migrate from the neural crest to the follicle, especially to the bulge, and further to the hair matrix and epidermis (14). *PAX3* is upregulated in melanoblasts derived from the neural crest, *KIT* is a prerequisite in melanocyte migration, and *DCT* is expressed in melanoblasts (15-17). The gene expression pattern indicates that the melanocytic differentiation of hESCs mimics the developmental process *in vitro*. Considering a stable gene expression of tissue-derived melanocytes *ex vivo*, hESC-derived melanocytes are able to contribute to future research on melanocytes and skin development.

The simple and straightforward protocol for successful melanocyte differentiation is probably due to the transient induction of JMJD3. JMJD3, an enzyme that demethylates H3K27me3, plays a vital role in the proliferation and differentiation of murine ESCs (18). JMJD3 is highly expressed in melanocytic nevi (19) and affects clonogenicity, self-renewal, and transendothelial migration in melanoma (20). In addition, JMJD3 is involved in the differentiation of keratinocytes in skin turnover (19). However, the contribution of JMJD3 to melanocytic differentiation has not been reported. JMJD3 activates BMP signaling (13), which is required for melanocytic differentiation (15). Likewise, transient expression of JMJD3 accelerates hepatic and myocytic differentiation (13). Given the involvement of JMJD3 in stem cell differentiation and oncogenesis, further research on JMJD3 in melanocytic differentiation will be of interest.

The long-time biocompatibility in mice of hESCs-derived melanocytes indicates that our protocol is useful in future clinical applications. Although vitiligo is not a life-threatening disease, 75% of patients with vitiligo feel distressed about their appearance and hesitate to go to new places due to their appearance (2). If the conservative treatment doesn’t work, a skin graft is the next option. However, adult melanocytes have the too low proliferative capacity *in vitro* to be used for treatment. Therefore, pluripotent stem cell-derived melanocytes have recently been the focus of much attention. hESC-derived melanocytes may not be engrafted due to their immunogenicity, especially compared to autologous cells derived from induced pluripotent stem cells (iPSCs). It is also noteworthy that hESCs-derived melanocytes are less likely to induce immunogenicity (21), and tissue-derived melanocytes indeed survived in immunocompetent mice. Direct subcutaneous injection of pluripotent stem cell-derived melanocytes can be used in the treatment of vitiligo since melanocytes injected into subcutaneous tissue migrate into the dermal-epidermal junction in mice (22), and can thus be a treatment due to its less invasiveness. ESCs with immortality can be used as raw material for on-the-shelf melanocyte preparations. The use of ESC-derived melanocytes for benign diseases such as vitiligo raises concerns about tumorigenesis and the benefits must be weighed against the risk.

## Conclusion

We report a simple and straightforward method to produce hESC-derived melanocytes with high proliferative ability over time, and functionality, simulating melanocytic development in embryogenesis. hESC-derived melanocytes share the characteristics of tissue melanocytes, and will thus be useful in clinical applications.

## Declarations

### Ethics approval and consent to participate

The institutional Review Board approved all experiments using human tissues at National Center for Child Health and Development. We obtained informed consent from all participants or their parents when participants were under 18. Human cells in this study were utilized in full compliance with the Ethical Guidelines for Medical and Health Research Involving Human Subjects (Ministry of Health, Labor, and Welfare (MHLW), Japan; Ministry of Education, Culture, Sports, Science and Technology (MEXT), Japan). The derivation and cultivation of human Embryonic Stem Cell (hESC) lines were performed in full compliance with “the Guidelines for Derivation and Distribution of hESCs (Notifications of MEXT, No. 156 of August 21, 2009; notifications of MEXT, No. 86 of May 20, 2010) and “the Guidelines for Utilization of Human Embryonic Stem Cells (notifications of MEXT, No. 157 of August 21, 2009; notifications of MEXT, No. 87 of May 20, 2010)”.

Animal experiments were performed in compliance with the basic guidelines for the conduct of animal experiments in implementing agencies under the jurisdiction of the Ministry of Health, Labour, and Welfare (notifications of MHLW, No. 0220-1 of February 20, 2015). The Institutional Animal Care and Use Committee of the National Center for Child Health and Development approved the protocols of the animal experiments. This study was carried out in compliance with the ARRIVE guidelines.

### Consent for publication

Not applicable.

### Availability of data and material

The datasets used and/or analyzed during the current study are available from the corresponding author upon reasonable request.

### List of Abbreviations

Not applicable.

### Authors’ contributions

CK, RT, RY, PJ, and AU designed experiments. CK, RT, PJ, and TA performed experiments. CK and RT analyzed data. SY, MM, HT, TA, and MSHK contributed reagents and materials. CK, RT, RY, PJ, TA, MSHK, KK, and AU discussed the data and manuscript. CK and AU wrote this manuscript. The authors read and approved the final manuscript.

## Acknowledgments

We would like to express our sincere thanks to M. Ichinose for providing expert technical assistance, to C. Ketcham for English editing and proofreading, and E. Suzuki and K. Saito for secretarial work.

